# GutMicroNet: an interactive platform for gut microbiome interaction exploration

**DOI:** 10.1101/2021.11.10.468051

**Authors:** Muhammad Arif, Theo Portlock, Cem Güngör, Elif Koç, Berkay Özcan, Oğuzhan Subaş, Buğra Çakmak, Hassan Turkez, Mathias Uhlén, Adil Mardinoglu, Saeed Shoaie

**Author notes:** To whom correspondence should be addressed., Correspondence may also be addressed to Adil Mardinoglu. The authors wish it to be known that, in their opinion, the first two authors should be regarded as Joint First Authors. Muhammad Arif, Laboratory of Cardiovascular Physiology and Tissue Injury and Section on Fibrotic Disorders, National Institute on Alcohol Abuse and Alcoholism, National Institutes of Health, Rockville, MD 20852, USA.

## Abstract

The human gut microbiome data has been proven to be a powerful tool to understand the human body in both health and disease conditions. However, understanding their complex interactions and impact on the human body remains a challenging task. Unravelling the species-level interactions could allow us to study the causality of the microbiome. Moreover, it could lead us to better understand the underlying mechanisms of complex diseases and, subsequently, the discovery of new therapeutic targets. Given these challenges and benefits, it has become evident that a freely accessible and centralized platform for presenting gut microbiome interaction is essential to untangle the complexity and open multiple new paths and opportunities in disease- and drug-related research. Here, we present GutMicroNet, an interactive visualization platform of human gut microbiome interaction networks. We generated 45 gut microbiome co-abundance networks from various geographical origins, gender, and diseases based on the data presented in the Human Gut Microbiome Atlas. This interactive platform includes more than 1900 gut microbiome species and allows users to query multiple species at the same time based on their interests and adjust it based on the statistical properties. Moreover, users can download publication-ready figures or network information for further analysis. The platform can be accessed freely on https://gutmicro.net without any login requirements or limitations, including access to the full networks data.

## INTRODUCTION

In recent years, the impact of the human gut microbial composition and their diversity on health and disease has become apparent. With the decreasing cost of metagenomic sequencing, availability and advances in high-performance computing, and the accessibility for researchers to collect sequencing data, analysis of metagenomic data has never been more achievable. Hence, gut microbial dysbiosis has been associated with the pathologies of a broad range of diseases, including obesity, inflammatory bowel disease and depression (1–3). Investigating the complex nature of the various interactions between bacterial communities in the gut and their impact on health and disease remains challenging. Unravelling the species-level interactions could allow us to study the causality of the microbiome and reveal the underlying molecular mechanisms involved in the progression of the disease. Previously, there have been different approaches in investigating the co-abundant species using different methods (4,5). However, none has focused on a systematic analysis of the species interactions across several cohorts and provided an interactive platform to explore the microbial networks.

Here, we developed GutMicroNet (https://gutmicro.net), an interactive platform to present the interactions between human gut microbiomes in health and diseases. In this platform, we presented microbial networks as species co-abundance networks based on the recent studies, where extensive gut microbiome data were collected from various geographical origins, gender, and diseases. Users can visualize multiple species, adjust the statistical properties, and interact with the visualized networks to explore interactions with other species, modify the network, and download the network as a static image or text file that can be imported to other network analysis software or modules for more complex downstream analyses. We envisage that GutMicroNet is a unique and powerful platform that will open multiple new paths and opportunities in disease- and microbiome-related research.

## MATERIAL AND METHODS

### Data Sources

The networks were constructed using the data presented in the Human Gut Microbiome Atlas (https://www.microbiomeatlas.org) (6). We retrieved the species abundance matrix and sample metadata tables from the atlas, and we further collected the metagenomic species pan-genomes (MSP) taxonomy(6) and KEGG Pathways (7) information from the atlas. The data contained 39 datasets, including 10 healthy from 9 countries (US, UK, Sweden, India, Fiji, Madagascar, Mongolia, Peru, and Tanzania), 29 diseases cohorts from 12 countries (US, China, Japan, UK, Sweden, Germany, Italy, Spain, France, Denmark, Finland, and Luxembourg). The details for each cohort can be found in **Supplementary Table 1**.

First, the data were split into each cohort and pre-processed independently. The species detected (abundance > 0) in less than 5 samples in a cohort were filtered. To generate the co-abundance network, the filtered data from the same geographical origin and/or disease (depending on the network type) were merged into a data-frame, analyzed using spearmanr function from SciPy package in a pairwise manner, and filtered to leave just the significant correlations (FDR < 0.05)(8). For the downstream analysis, we performed community detection using the Leiden algorithm(9), centrality analysis, and cluster average local transitivity (clustering coefficient) from the iGraph Python (10) package and functional analysis with enrichr function in the GSEAPY package(11–14) using the MSP-to-KEGG mapping from the atlas. All analyses were performed in Python 3.7.4.

### Platform Description and Features

The GutMicroNet (https://gutmicro.net) contains the interactions of 1976 gut microbiome species from various geographical origins and health conditions stored in the Neo4j graph database. There are two ways of accessing the platform: (1) a web-based platform for all users, including scientists with no bioinformatics background, and (2) programmatic access for more advanced users. Moreover, the selected networks can also be downloaded as publication-ready figures (only via web) or as network tables (both web and programmatic access) that can be integrated directly with other network-specific software, e.g. Cytoscape(15).

### Web Platform Interactive Access

The primary way to access the platform is via an interactive, intuitive, and user-friendly web interface on https://gutmicro.net (**Figure 1A-B**). On this website, users can visualize their network of interest with 2 simple steps: (1) choosing the specific network category and network type (based on geography or diseases) and then, (2) query for up to 5 species. The web platform provides the possibility to flexibly move the nodes and trim or expand their network, after the initial visualization, by (1) adding more species in the query, (2) clicking on the “ADD” or “REMOVE” button on the node information panel and (3) adjusting the statistical variables, i.e. FDR and co-abundance rank (based on absolute correlation score). The users can also choose the taxonomic level to group the nodes by, represented by nodes colours. The node colour codes will be updated dynamically and included in the network legends based on users’ choices. Once the users are satisfied with their networks, they can be exported as ready-to-use figures or tables (including the node colour codes) to be analysed further with other network analysis software.

**Figure 1:**
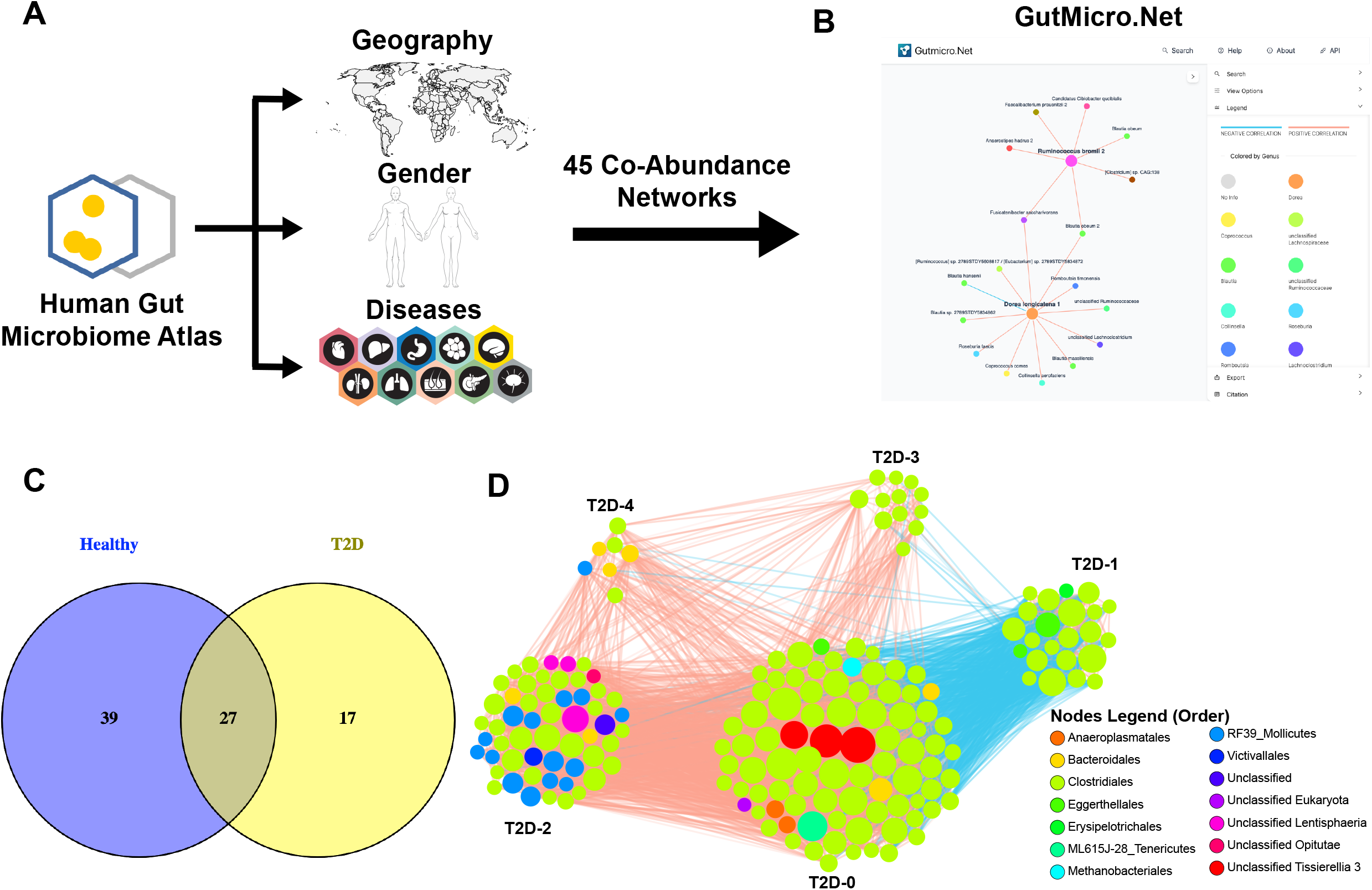
**(A)** The primary data source of GutMicroNet was data from various geographical origins, gender, and diseases originated from the Human Gut Microbiome Atlas **(B)** Interactive web access interface of GutMicroNet **(C)** Venn Diagram showing the similarities and differences of the central nodes of Healthy and T2D network **(D)** Communities detected on the T2D network (including only edges with FDR < 0.001 and nodes with degree centrality > 50)

### Programmatic Access

GutMicroNet can also be accessed programmatically using in-house Python code. Under the “API” section on the website, we have provided the Jupyter Notebook with an example of accessing GutMicroNet programmatically and how to integrate it with the iGraph network tool. With programmatic access, more advanced users can automatize the query, incorporate the networks into their data/analyses, retrieve more complex networks, and perform more comprehensive downstream analysis based on their needs.

## RESULTS

Based on the Human Gut Microbiome Atlas (6) data (**Figure 1A**), we generated and included 45 MSP co-abundance networks, including overall, geography-, gender-, and disease-specific networks, to GutMicroNet with various network sizes. We observed high variance in the network size, which could be due to the nature of microbial abundance data and the sparsity and high variation, together with the cohort size (**Supplementary Table 2**). The number of edges and nodes varied from 1101 to 568.008 edges and 355 to 1.929 nodes, with colorectal cancer from Italy and the overall network showing the lowest and highest density, respectively. Several networks were discarded due to the low number of samples and interactions.

In GutMicroNet, users can choose the networks that fit their studies, from global networks to gender-, geography-, and disease-specific networks **(Figure 1B).** Moreover, GutMicroNet has this unique capability to generate co-abundance networks at a personalized level, which means that individual variations and biases were eliminated. In this section, we presented two case studies to elucidate the strength of GutMicroNet in microbiome-related research. In the first case, we used the centrality and community detection of the downstream network analyses to identify the changes in coabundance network structure in cohorts of type-2 diabetes. In the second case study, we showed how GutMicroNet could be used to validate and discover new insights based on the preliminary findings in our previous NAFLD-related research.

### Case Study 1: Comparing Healthy and Disease Microbiome-Interaction Networks

GutMicroNet provides a vast number of microbiome interaction networks, including healthy and disease networks from different populations. This feature makes it possible for us to analyze the changes in biological network structure that have been known to be associated with the changes in pathological conditions (16). In this case study, we compared the core network structure of healthy and disease networks from the same geographical origin (Sweden), represented by “Wellness Cohort” (17) (subsequently referred to as Healthy network) and “Type-2 Diabetes (Sweden)” (T2D network), to identify the changes in the top connected nodes in each network. Moreover, we attempted to identify the most critical network community in both conditions and their functional property.

*Key Bacterial Family Differs between Healthy and T2D Networks.* There are many approaches to define the most or critical nodes in a network, usually referred to as centrality analysis, such as betweenness and closeness centrality. In this case study, we used the most common centrality measure: degree centrality. Degree centrality represents the number of directly connected neighbours to a node. Centrality analysis, specifically with degree centrality, has been proven to help scientists in disease-related research, such as drug repositioning(18).

We calculated the degree from both networks and filtered for the top 5% of most connected nodes (66 and 44 nodes in the healthy and T2D network). From those nodes, we found that 27 nodes were shared in both networks (**Figure 1C**). This shows that similarities still could be detected in the samples from the same geographical origin regardless of the health conditions. One of the standouts shared central nodes is *Blautia wexlerae,* as they were found to be enriched in the Swedish population and both healthy and T2D samples, based on the Human Gut Microbiome Atlas (https://www.microbiomeatlas.org/species.php?species_msp=msp_0076).

We also found several differences in the central nodes. Noticeably, four bacteria families were eliminated as central nodes in the T2D: *Bacteroidaceae (Bacteroides vulgatus)* (19), *Methanobacteriaceae (Methanobrevibacter smithii 1)* (20), *Atopobiaceae (Olsenella sp. GAM18)* and *unclassified RF39 Mollicutes.* Interestingly, based on the literature, the alteration of *Bacteroides vulgatus* and *Methanobrevibacter smithii 1* have also been associated with the pathology of T2D (21). Moreover, 2 bacterial families were introduced as the key nodes in T2D: *Odoribacteraceae (Odoribacter splanchnicus)* and *unclassified Clostridiales 1. Odoribacter splanchnicus* was shown as a depleted species in Type 2 diabetes (https://www.microbiomeatlas.org/species.php?species_msp=msp_0062) and were previously associated with lipid profile in obesity (22).

*Community Detections.* Biological networks, including co-abundance gut microbiome networks in our platform, are generally complex and large. To understand a network properly, we need to disentangle its complexity. One way to do that is by performing community detections to split a large network into several smaller subnetworks with high connectivity within them. These subnetworks are usually referred to as communities or clusters. Community detections assisted researchers in getting hidden insights into biological networks, such as in the multi-tissue gene co-expression networks of heart disease (23) or integrated networks in NAFLD (24).

We performed the Leiden community detection algorithm in both the healthy and T2D networks. We found 5 clusters with > 30 nodes in both networks (**Figure 1D**). Subsequently, we identified the most driver community in both networks by selecting the communities with the highest average clustering coefficient (24): Healthy-4 (80 nodes) and T2D-3 (149 nodes). On the functional level, we found that the members of the central clusters were associated with several important functions as glycolysis, lipoic acid metabolism, and amino acid metabolism (histidine; cysteine and methionine; phenylalanine, tyrosine, and tryptophan biosynthesis). Specifically, the species in Healthy-4 were also associated with vitamin B6 metabolism, TCA cycle, and lipopolysaccharide biosynthesis. On the other hand, the species in T2D-3 were related to known T2D related pathways, such as pyruvate metabolism (25), galactose metabolism (26), and glycine, serine, and threonine metabolism (27).

Furthermore, we observed that Healthy-4 (80 nodes) was composed of species from *Lachnospiraceae* (8 species) and *Ersipelotrichaceae* (7) families; meanwhile, T2D-3 (149) was dominated by *Lachnospiraceae* (25) and *Ruminococcaceae* (21) families. Interestingly, even though *Lachnospiraceae* was abundant in both central clusters, the genus (and species) were totally different in Healthy-4 *(Blautia, Eisenbergiella,* and *Roseburia)* and T2D-3 *(Dorea, Faecalicatena, Lachnoclostridium, Marvinbryantia, Merdimonas, Mordavella,* and *Tyzzerella),* except the *Lachnoclostridium* and *unclassified Lachnospiraceae genus.* Moreover, we found several other families that were previously associated with T2D in the T2D-3 cluster, e.g. *Akkermansiaceae (Akkermansia muciniphila 2)* (19,28,29) and *Clostridiaceae (Clostridium disporicum)* (30).

The first use case showed the strengths of this platform to explore the dysbiosis of the gut microbiome in metabolic-related complex diseases by comparing the network structures between healthy and disease-specific networks. These approaches can be placed as pre-study exploratory analysis or to complement the biological experiment results. Even though we focused on the known and validated species interactions and functions in this use case, we also found many new unknown interactions that can open new avenues in understanding the disease and, possibly, new therapeutic targets.

### Case Study 2: Validating the gut microbiome dysbiosis in NAFLD

Not only that GutMicroNet includes a large number of networks, but the networks in the platform were also generated from personalized microbiome data. This resulted in the minimization, if not elimination, of bias caused by individual variations, a known problem in microbiome data analysis. In this case study, we attempted to use GutMicroNet to validate our findings in our recent research on the association of microbiota and non-alcoholic fatty liver disease (NAFLD) (31).

In that study, we collected multi-omics data, including gut metagenomics data, from 56 individuals with NAFLD and varying levels of hepatosteatosis (HS). The samples were stratified based on their proton density fat fraction measured by magnetic resonance imaging (MRI-PDFF) results. Subsequently, we performed differential expression, correlation network, and random forest analysis to determine omics signatures of each HS group. Among those critical signatures, we found two gut microbiota species to be negatively associated with liver fat and enzymes, and good predictors of HS: *Dorea longicatena* and *Ruminococcus bromii.* Using GutMicroNet, we were able to validate the association of those species with NAFLD. We retrieved the neighbour of *Dorea longicatena* and *Ruminococcus bromii* in the “NAFLD Non-Diabetic” network (**Figure 2**). Interestingly, we found 5 neighbouring species that were found to be associated with NAFLD by the Human Gut Microbiome Atlas: *Blautia obeum, Blautia obeum 2, Blautia sp. 2789STDY5834862, Collinsella aerofaciens,* and *Faecalibacterium prausnitzii 2.* Moreover, both species shared a neighbour, *Fusicatenibacter saccharivorans,* that has been shown to be negatively associated with liver fat (32). Furthermore, we checked the functional association of the network nodes and found that they were related to several NAFLD-related KEGG pathways, including fatty acid biosynthesis, serine, cysteine, glycine, and tryptophan metabolism.

**Figure 2:**
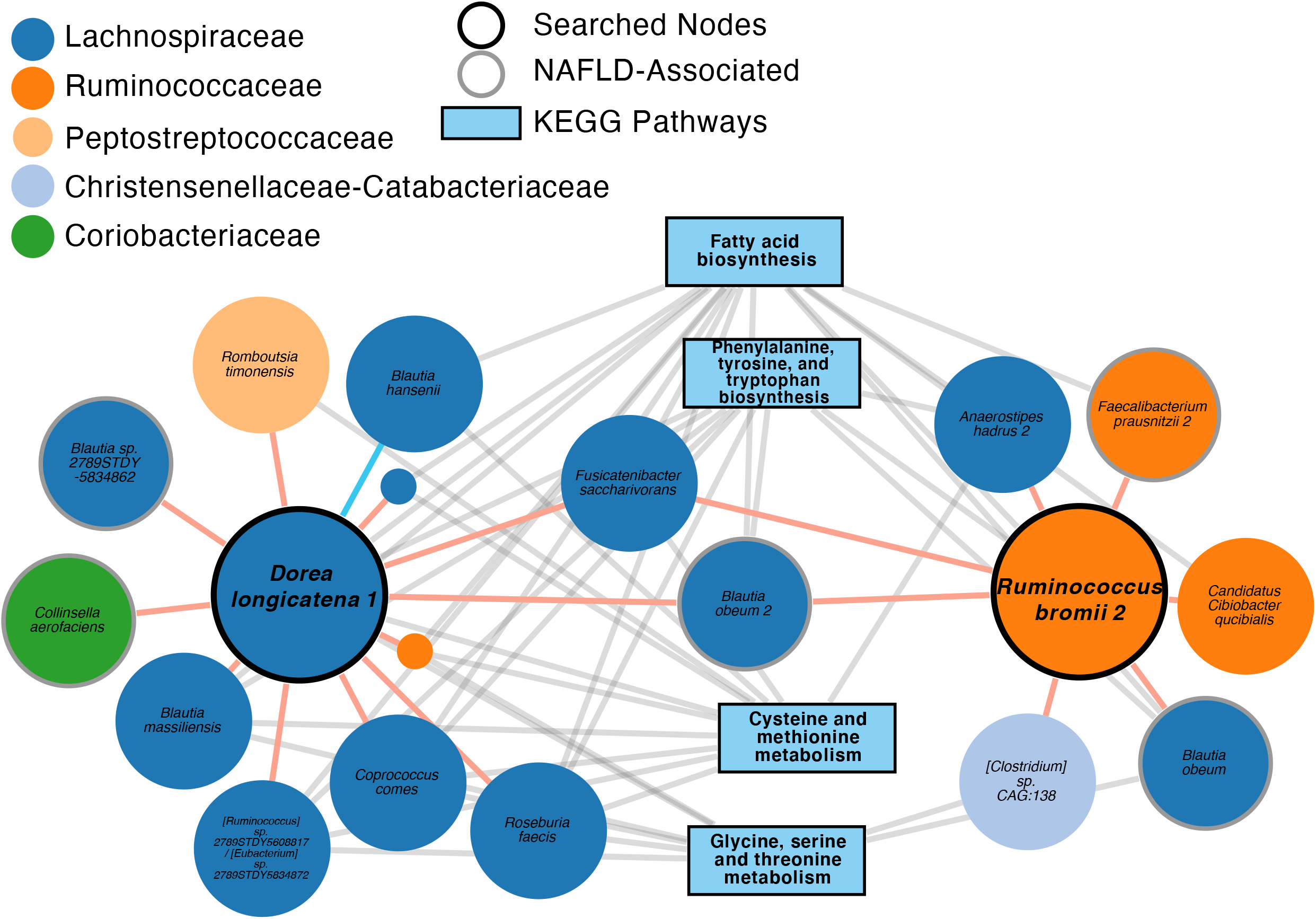
Validation of an independent novel non-invasive NAFLD biomarkers, *Dorea longicatena* and *Ruminococcus bromii,* using the NAFLD Non-Diabetic network from GutMicroNet.

In this case study, we showed how GutMicroNet could be employed as an in-silico validation tool in biomarker-discovery research. Not only that it strengthens the findings, but also shortens the analysis cycle since it is freely and easily accessible.

## DISCUSSION

In this study, we introduce GutMicroNet (https://gutmicro.net), an interactive web-based platform to explore the interactions of gut microbiota species. The interactions are presented in 45 co-abundance networks, including overall and gender-, geography-, and disease-specific networks, based on data from Human Gut Microbiome Atlas (https://www.microbiomeatlas.org/). As shown in both case studies, this platform provides multiple features that may assist scientists in performing pre-study exploratory analysis, identifying features and functions associated with the microbes, and validating their experiment results. Moreover, this platform offers the possibility of discovering new and hidden insights that may lead to novel biomarkers and therapeutic targets. On top of the features, we built this platform with user experience in mind, which resulted in an intuitive and user-friendly design for users regardless of their bioinformatics or systems biology background.

We foresee that GutMicroNet will be an essential tool for researchers and scientists, especially those working with the gut microbiome. We also aim to improve this platform and make it the standard and go-to resource for microbiome research. In the next version of GutMicroNet, we plan to expand the platform by adding more studies to improve the quality and variation of the networks. Furthermore, we plan to add a feature to personalize the pre-generated networks by integrating users’ experimental data into our platform.

## DATA AVAILABILITY

All network data can be found on https://gutmicro.net. MSP tables and metadata can be found on the Human Gut Microbiome Atlas download section (https://www.microbiomeatlas.org/downloads.php)

## SUPPLEMENTARY DATA

Supplementary Data are available at NAR online.

---

**Supplementary Table 1:** Detailed cohort information from the Human Gut Microbiome Atlas

**Supplementary Table 2:** Complete network information

